# Wheat stripe rust resistance gene *Yr9*, derived from rye, is a *CC-NBS-LRR* gene in a highly conserved *NLR* cluster

**DOI:** 10.1101/2024.10.04.616745

**Authors:** Yang Yu, Jiajun Liu, Shengjie Lan, Qihang Chen, Jinlong Li, Haoyuan Song, Chen Pan, Juan Qi, Fei Ni, Lynn Epstein, Daolin Fu, Jiajie Wu

## Abstract

Wheat stripe rust, caused by *Puccinia striiformis* f. sp. *tritici* (*Pst*), is a significant threat to global wheat production. Genetic resistance plays a crucial role in controlling this disease. Among wheat breeding innovations, the wheat-rye 1BL.1RS translocations are notable for introducing alien genetic diversity, partly due to the presence of the stripe rust resistance gene *Yr9* on 1RS. To clone *Yr9*, we first identified four *Pst*-susceptible mutants from Lumai 15, which carries the 1BL.1RS translocation and *Yr9*. Using these mutants, we performed Sequencing Trait-Associated Mutations (STAM). A single candidate gene, *YrChr1B*, was identified within the *Yr9* locus and later confirmed as *Yr9* through genetic complementation and gene editing. The *Yr9* gene encodes a coiled-coil nucleotide-binding site leucine-rich repeat (CC-NBS-LRR or NLR) protein and is part of a 14-member *NLR* gene cluster. This cluster is conserved among Triticeae species and is an ortholog of the barley *Mla* locus. Cloning *Yr9* expands the genetic resources available for molecular wheat breeding aimed at durable and broad-spectrum disease resistance.

## Introduction

Crop pathogens cause substantial economic losses and reduce food security globally (1). Harnessing disease resistance genes and breeding resistant crops are the most effective ways to reduce losses caused by pathogens. However, domestication and cultivar breeding of wheat and other crops have significantly reduced their genetic diversity (2-3). The reduced diversity has resulted in a shortage of disease resistance genes, which is a severe obstacle for wheat breeding. Wheat relatives are valuable sources of genetic diversity that can improve wheat’s disease resistance. In wheat breeding history, the replacement of the short arm of chromosome 1B from wheat with the 1RS from rye (*Secale cereale* L.), has been the most impactful incorporation of alien genetic diversity (4). The 1BL.1RS translocation enhances wheat resistance to biotic and abiotic stresses, and provides resistance to stripe rust (yellow rust, *Yr9*), leaf rust (*Lr26*) and stem rust (*Sr31*). Previous studies revealed that *Yr9, Lr26* and *Sr31* are separate, but closely-linked genes that co-segregate with *Mla-LRR* markers (5). However, attempts to clone these genes have been unsuccessful. Despite significant global efforts to clone stripe rust resistance genes in wheat, only ten genes for resistance to *Pst* have been successfully cloned to date (6). Although *Yr9, Lr26*, and *Sr31* have been defeated by newly emergent virulent races, revealing the resistance locus on 1RS will facilitate mining untapped genetic diversity and provide new insights into the evolution of plant resistance genes. To address this question, we show that *Yr9* encodes a *NLR* gene, is in a R gene cluster that is conserved among *Triticeae* species and is an ortholog of the barley *Mla* locus (7).

## Results

‘Lumai 15’ (LM15; *Triticum aestivum* L.), a predominant Chinese wheat cultivar released in the 1990s, is a 1BL.1RS derivative that yields well and is disease resistant (8). Because of the 1RS translocation, when LM15 is challenged with *Pst* races CYR23 and CYR34, seedling leaves are immune (infection types IT=0) (Fig. 1*A*). Another 1BL.1RS cultivar ‘Aikang 58’ (AK58) shows slight chlorotic and necrotic flecks with a resistant infection type (IT=1) (Fig. 1*A*), indicating that the enhanced resistance of LM15 might be due to additive effect of *Yr9* and other unknown resistance gene(s).

**Fig. 1.**
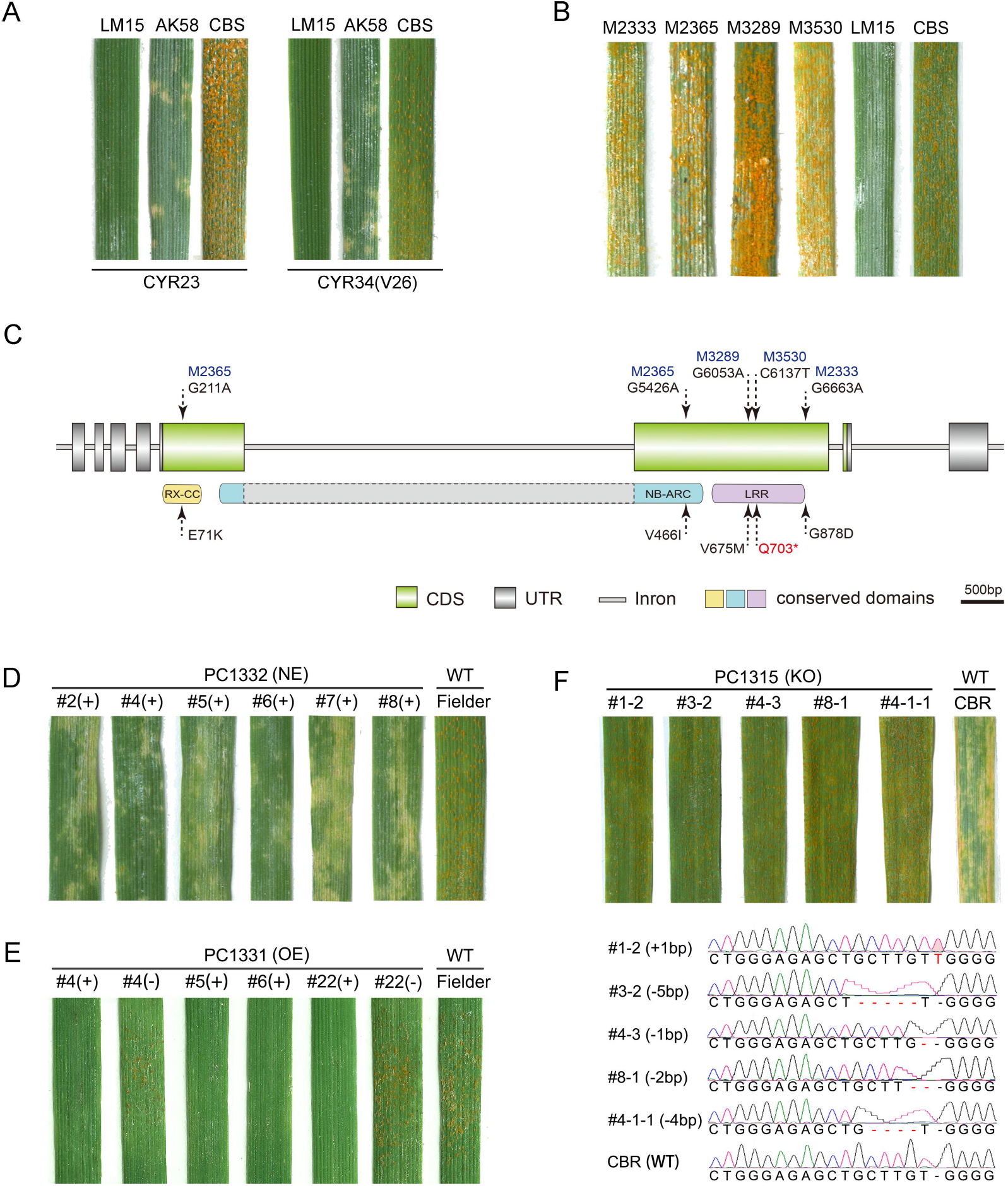
The cloning and functional validation of *Pst*-resistance gene *Yr9*. (*A*) Phenotypic responses of 1BL.1RS cultivars Lumai 15 (LM15) and Aikang 58 (AK58) to *Pst* races CYR23 and CYR34 (V26), with CB037-*Pst*S (CBS) as the susceptible control. (*B*) Four *Pst*-susceptible mutants of LM15. (*C*) The *YrChr1B* (*Yr9*) gene and conserved domains. Mutations and amino acid changes are labeled. (*D*) Responses of six independent T_1_ transgenic lines transformed with natural-expression (NE) vector PC1332, with WT Fielder used as the susceptible control. (*E*) Responses of T_1_ transgenic lines derived from the over-expression (OE) vector PC1331. (*F*) CRISPR/Cas9 editing of *Yr9* in 1BL.1RS cultivar CB037-*Pst*R (CBR). WT CBR is resistant to CYR34, and *Yr9* edited plants are susceptible. Small insertions or deletions occurred at the sgRNA target of T_2_ (#4-1-1) and T_1_ (four other) plants. Symbols ‘(+)’ and ‘(−)’ in *D* and/or *E* represent positive and negative siblings, respectively.

To identify the *Pst*-resistance genes in LM15, an ethyl-methanesulfonate (EMS) mutagenized population was propagated and used to screen for *Pst*-susceptible mutants. Among the 2,800 M_3_ generation lines, individual plants from four lines (M2333, M2365, M3289, and M3530), were identified as highly susceptible (ITs=7-9) (Fig. 1*B*) and propagated. In M_5_ generation, the most susceptible plant was selected from each line, and then used for Sequencing Trait-Associated Mutations (STAM) analysis (9).

In STAM analysis, the Iso-seq from wild-type (WT) LM15 produced 55,716 non-redundant isoforms that were used as a reference (*Dataset S1*). RNA-Seq reads from the four mutants were used to call SNPs between the WT LM15 and *Pst*-susceptible mutants. Only high-confidence SNPs that were homozygous, and EMS-prone mutations (G to A, or C to T), were analyzed. A total of 2,083 transcripts that have SNP(s) in at least one mutant line were identified. Only one transcript (#1460) (GenBank PQ412819) was mutated in all four lines; M2365 has two missense mutations, M3530 has one nonsense mutation, and M2333 and M3289 each has a missense mutation (Fig. 1*C*). We named the transcript #1460 as *YrChr1B*; its coding region is 2,889 bp and encodes an iconic, intracellular CC-NBS-LRR disease resistance gene (Fig. 1*C*).

In Blastn searches against assembled wheat genomes, *YrChr1B* is 100% identical to TraesKN1B01HG01710.1 (located at 15.47 Mb) in 1BL.1RS cultivar ‘Kenong 9204’ (KN9204) (10), and has 99% similarity with SECCE1Rv1G0003760.1 (located at 14.97 Mb) in rye ‘Lo7’ (11). In comparison, the most similar genes to *YrChr1B* in the reference ‘Chinese Spring’ are located in 1DS and 1AS with sequence identities of 89% (TraesCS1D03G0054800.1) and 92% (TraesCS1A03G0066200.1). We further found that the physical position of *YrChr1B* is rather close to homologues of the RFLP marker BE196644 (GenBank) that co-segregates with *Yr9* (5). This barley EST BE196644 (448bp) is located at 15.25 Mb of 1BS (1RS) of KN9204 and 14.92 Mb of 1RS of Lo7.

To determine if *YrChr1B* confers *Pst-resistance*, we constructed PC1332, a natural expression vector with 9.3 kb genomic sequence (7,053 bp from the start to stop codons plus 2,322bp upstream). After transforming *Pst-*susceptible cultivar ‘Fielder’ with PC1332, we showed that *YrChr1B* significantly enhances *Pst*-resistance (Fig. 1*D*). The over-expression of *YrChr1B* coding sequence (PC1331) also confers *Pst*-resistance (Fig. 1*E*). *YrChr1B* function also was confirmed with CRISPR/Cas9 gene-edited plants. Using a 1BL.1RS cultivar ‘CB037-PstR’ (CBR) as WT, *YrChr1B* knock-out plants were obtained by transformation with PC1315, which expresses sgRNAs that target the 5^th^ exon of *YrChr1B*. Whereas WT CBR is highly resistant, *YrChr1B-* edited plants are highly susceptible (Fig. 1*F*). Thus, *YrChr1B* confers *Pst-*resistance in 1BL.1RS cultivars LM15 and CBR. Given that only one *Pst*-resistance gene (*Yr9*) was mapped to 1RS (5), *YrChr1B* is *Yr9. Yr9* expression exhibits a marked decrease, reaching its lowest level 24 hours post-*Pst* inoculation in WT LM15, CBR, and PC1332 transgenic (*Dataset S2*).

The four 1BL.1RS cultivars (LM15, KN9204, CBR, and AK58) have an identical *Yr9* sequence. In KN9204, *Yr9* is within a *NLR* gene cluster, flanked by TraesKN1B01HG01640 (benzyl alcohol O-benzoyltransferase-like, BEBT) and TraesKN1B01HG01740 (nucleic acid-binding protein, NABP) (Fig. 2). In this 430 kb region, eight *NLRs* were annotated in a whole genome sequencing project (10). By sequence alignment and structure prediction, we annotated six more *NLRs* in this cluster (*Dataset S3*). Based on our Iso-Seq transcripts, five of 14 *NLRs* are transcribed (Fig. 2). This *NLR* gene cluster is conserved among *Triticeae* species; alongside the conserved *NABP* gene (Fig. 2). Interestingly, this region is orthologous to the barley *Mla* locus that confers powdery mildew resistance (7). Stem rust resistance gene *Sr50*, from *S. cereale* cv. ‘Imperial’, is also in this orthologous region (12). Therefore, diverse resistance genes to fungal diseases have evolved in the *Yr9* ortholog region. In addition, chymotrypsin inhibitor 2 (*CI2*), which is another type of gene that is correlated with plants’ resistance to wounding and pests (13) is also enriched around the *Yr9* locus (Fig. 2).

**Fig. 2.**
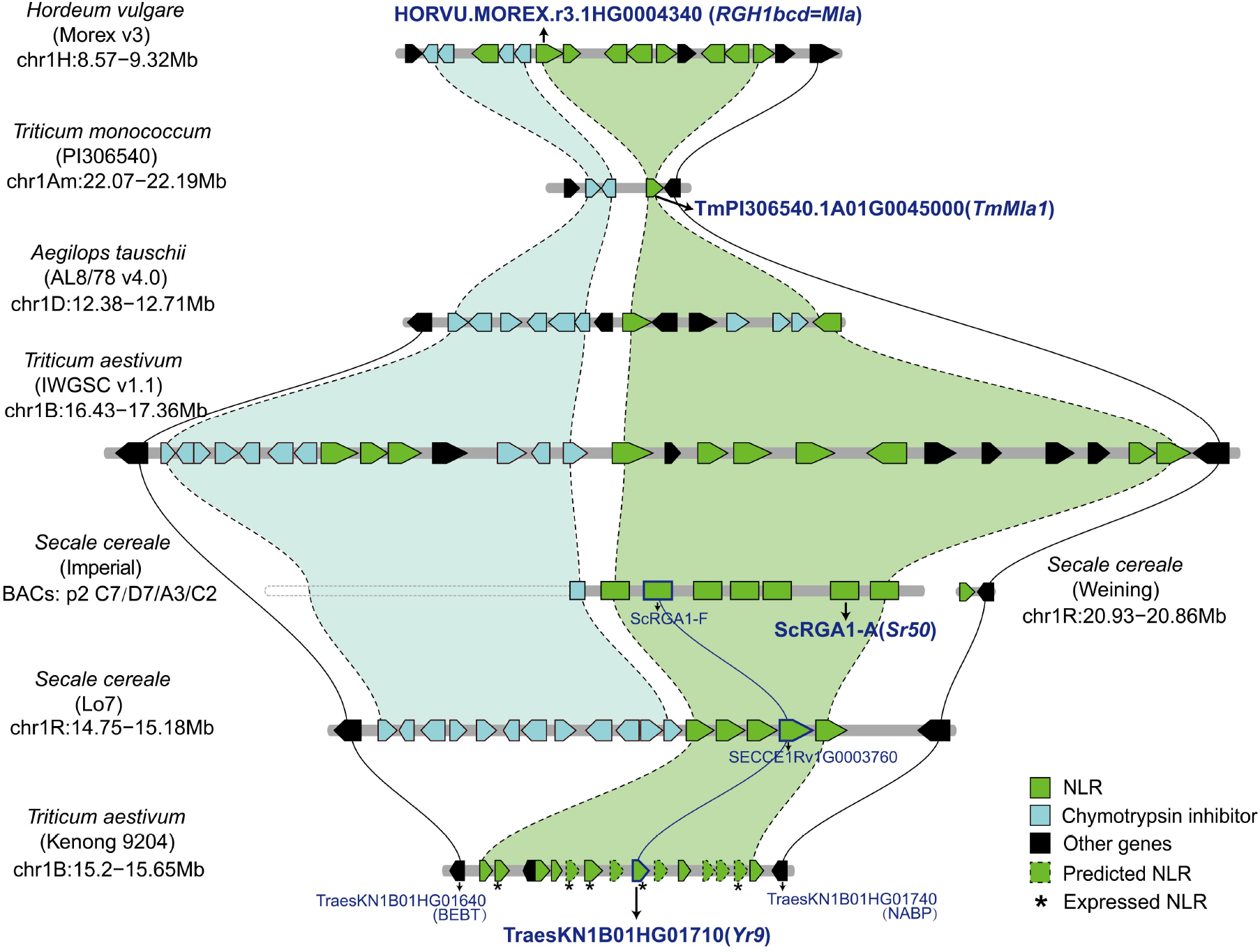
The conserved *NLR* gene cluster harboring *Yr9*. Using the *Triticeae*-Gene Tribe tool, we modified the output to show the collinearity between *T*.*aestivum* cv. KN9204 (1BL.1RS derivative) and Chinese spring, *T*.*monococcum* acc. PI306540, 1R of *S*.*cereale* cv. Lo7 and ‘Weining’, 1D of *Ae*.*tauschii* acc. ‘AL8/78’, and 1H of *H*.*vulgare* cv. ‘Morex’. The physical map of *Sr50* from *S. cereale* cv. Imperial was added manually. In KN9204, six additional *NLR* genes were annotated manually and highlighted with dashed boxes; *NLRs* with confirmed expression are indicated by an asterisk.

We further investigated the distribution of *Yr9* in rye germplasm. Using two primer pairs, *Yr9* was amplified from 6 of 19 lines (*Dataset S4*). We also analyzed public re-sequencing data of rye Lo7, Weining and another 13 inbred lines (11,14-15). The results showed *Yr9* is present in Lo117, Lo282, Lo298 and Lo348; *Yr9* might also be present in Lo351, R925 and R1003, but absent in Weining, R2446 and other 5 inbred lines (*Dataset S5*).

## Discussion

We show that *Yr9*, which functions in rye and in wheat 1BL.1RS derivatives, encodes for a NLR protein that is located in a *NLR* cluster. Consequently, it would have been difficult to identify *Yr9* through traditional map-based cloning. Here we cloned *Yr9* using STAM (9) with only four EMS-prone mutants, instead of seven or more independent mutants used previously for STAM analyses or similar strategies (16).

The *Yr9*-associated *NLR* cluster is conserved among wheat, rye, and barley. It is an ortholog of the *Sr50* rye locus (12), the barley *Mla* locus (7), and the *monococcum* wheat *TmMla1* locus (17). Therefore, this region contains genes that confer resistance to multiple pathogens. *TmMla1* provides resistance to powdery mildew in both wheat and barley (17). Based on PCR amplification and genome analysis, *Yr9* is present in approximately 30% of rye germplasm (*Dataset S4*&*5*). Exploration of the genetic diversity of this region among more species and cultivars will help identify more functional resistance genes and will provide more insights into the evolution of *NLR* clusters.

## Materials and Methods

### EMS mutagenesis

The 1BL.1RS wheat cultivar Lumai 15 (LM15) exhibits resistance to *Puccinia striiformis* f. sp. *tritici* (*Pst*), particularly against tested races such as CYR23, CYR34 (V26), as well as natural spores collected in Tai’an, Shandong Province. The wild-type LM15 seeds were treated with EMS (Sigma-Aldrich Co., St. Louis, MO, USA) following the protocol outlined in Ni *et al*. (2023) (9). Approximately 2,800 M_3_ mutant lines were harvested in the summer of 2018. This EMS population was screened for *Pst-*susceptible lines in greenhouse (1,700 lines) and field trials (1,100 lines). Six seeds from each line were planted in either a single pot in the greenhouse or a single row in the field. For *Pst* inoculation, a 2.5mL syringe was used to inject an aqueous spore suspension into the leaf bundles at the seedling stage, with three sequential injections at 7 d intervals. The *Pst* spores were natural mixtures collected from susceptible plants grown at the field station of Shandong Agricultural University. Infection types (ITs) were recorded 25 days post inoculation using a 0-9 scale, where ITs 0-3 were considered resistant, 4-6 intermediate, and 7-9 susceptible (18). Segregation of susceptible individuals was observed in four lines during the M_3_ generation. Three of these lines (M2365, M3289, and M3530) were identified in the greenhouse and the remaining line (M2333) was identified from the field. In the M_4_ and M_5_ generations, all lines, except for line M2333, continued to segregate for resistance and susceptibility.

### Sequencing trait-associated mutations (STAM)

To perform STAM analysis, first, Iso-Seq and RNA-Seq were performed on the wild-type (WT) LM15 and four *Pst-*susceptible mutants, respectively. Leaves from WT LM15 and four EMS mutants were collected after phenotypic verification through *Pst* inoculation. Total RNA from inoculated leaves was extracted using TRIzol reagent (Tiangen Biotech Co., Beijing, China) to construct cDNA libraries. Sequencing of LM15 was performed using the PacBio Sequel II system (Pacific Biosciences, CA, USA), while the EMS mutants were sequenced on the Illumina platform (2 × 150 bp paired-end high-throughput sequencing) by Berry Genomics (Beijing, China). The WT LM15 sequencing produced 205,035,786,930 subread bases with a subread N50 of 1,766 bp, while each EMS mutant generated approximately 14 Gb of raw sequencing data. Raw sequencing data have been deposited in the NCBI Sequence Read Archive (SRA) under accession number PRJNA1166628.

STAM analysis was conducted as described in Ni *et al*. (2023) (9). Briefly, *de novo* transcriptome reconstruction was performed using ccs (Version 6.4.0), lima (Version 2.2.0), and IsoSeq3 (Version 3.8.0). To filter out *Pst* gene contamination, repetitive DNA, and redundant isoforms, tools such as Minimap2 (v2.24), RepeatMasker (http://www.repeatmasker.org), pbmm2 (https://github.com/PacificBiosciences/pbmm2), Cogent (https://github.com/Magdoll/Cogent), and cDNA_Cupcake (https://github.com/Magdoll/cDNA_Cupcake) were used for data cleaning. A total of 98,212 full-length non-concatemer (FLNC) isoforms were assembled. Redundancy was reduced using CD-HIT-EST with a 0.99 similarity threshold, resulting in 55,716 non-redundant transcripts. RNA-seq reads from the four mutants were mapped to these non-redundant reference transcripts using STAR (v2.7.10a). SNP calling was performed using GATK v4.2, and preliminary variant filtering was applied with the following parameters: “DP < 5 || FS > 60.0 || MQ < 40.0 || QD < 2.0”. Further SNP screening criteria included: 1) read depth ≥ 5, 2) homozygous mutations, 3) EMS-prone mutations (G to A and C to T transitions), and 4) mutations unique to one sample. In the previous study (9), we retained SNPs that were detectable in >60% (four out of seven) of the mutants. In this study, we adjusted this parameter to ≥ 50% (two out of four). Initially, one SNP per transcript was permitted for each mutant. However, considering that some genes are quite long and may harbor more than one mutation, up to two mutation sites per transcript were allowed within a single sample.

In this study, a total of 2,083 transcripts that have SNP(s) in at least one mutant line were identified. Of those, only transcript #1460 had a mutation in all four mutant lines. The presence of four mutants was necessary because, by coincidence, some transcripts had mutations in multiple mutants: three of the lines had a mutation in two transcripts, two of the lines had a mutation in 70 transcripts, and 2010 transcripts had a mutation in only one of the lines.

Accurate phenotyping of the mutants is the most critical step in STAM. Here, there were only four highly susceptible mutants identified in 2,800 LM15 M_3_ lines. This ratio is very low compared to similar studies. This discrepancy is partly due to the presence of other *Pst* resistance gene(s) in LM15. In the four *Pst*-susceptible mutants, there may be other resistance gene(s) that are also mutated, and are either homozygous or heterozygous in those resistance alleles. Nevertheless, all three functional validation experiments confirmed that *YrChr1B* confers resistance, and is a wheat R gene.

### Functional validation methods

Natural-expression (NE), over-expression (OE), and CRIPSR/Cas9 based knock-out (KO) vectors were constructed for *Yr9*. For the NE construct, the full-length of *Yr9*, along with 2,322 bp upstream of the start codon (a total of 9,375 bp), was amplified from a BAC (Bacterial Artificial Chromosome) clone P978 which was screened from the BAC library of LM15_*RMs2*_ (19). The PCR amplification was carried out using high-fidelity DNA Polymerase Atelta Max (Vazyme Biotech, Nanjing, China), with a forward primer of 5’-GCTCTTAGCGTTTCTCTTTTGTAGTAGCT-3’ and a reverse primer of 5’-TCAGATCTCCTCGTCCTCGCAC-3’. The *Yr9* cassette was initially inserted into a pENTR backbone using pBM 27 toposmart cloning kit (Biomed, Beijing, China), and then transferred into plasmid PC613 (maintained in our lab) using LR Clonase™ II (Invitrogen, Waltham, USA). The final NE construct was designated PC1332.

For the OE construct, a 2,889 bp fragment, containing only the coding sequence from the start to the stop codons, was amplified from LM15 cDNA with forward primer 5’-ATGGATATTGTCACGGGTGCCA-3’ and reverse primer 5’-TCAGATCTCCTCGTCCTCGCAC-3’, and cloned into plasmid PC186, resulting in the final construct PC1331.

For the KO construct, two sgRNA sequences, 5’-CTGGGAGAGCTGCTTGTGG-3’ and 5’-ATTCAAGGGGCTCGTCAAG-3’, targeting 43-61 bp and 294-312 bp downstream of the start codon of *YrChr1B*, respectively, were inserted into the pBUE413 vector backbone. The pBUE413 plasmid, along with the intermediate cloning vector pCBC-MT1T2, were provided by Prof. Zhongfu Ni from China Agricultural University. The final KO construct was designated PC1315.

All final constructs were transformed into *Agrobacterium* strain EHA105. Genetic transformation of common wheat was performed using the *Agrobacterium*-mediated transfer method (20). The wheat cultivar ‘Fielder’, which is susceptible to stripe rust, was used as the recipient for PC1331 and PC1332, while the 1BL.1RS cultivar CB037-PstR (CBR) (21) served as the recipient for PC1315.

Positive transgenic plants were verified through PCR amplification of the *Yr9* gene for PC1331 and PC1332 transgenics, while sequencing of the sgRNA-targeted region was used for PC1315 transgenics. T_0_, T_1_ and T_2_ lines were tested for resistance to *Pst* at the seedling stage by applying a mixture of spores and talc (or *Lycopodium*) powder onto the leaf surface, as described by Ni *et al*. (2023) (9). *Pst* race CYR34 (V26), provided by Prof. Jianhui Wu from the Northwest A&F University, were used in this study.

### Gene expression analysis

At the two-leaf stage, WT LM15, CBR, and PC1332 (NE construct) transgenics were inoculated with *Pst* race CYR34 (V26) under controlled conditions (22°C day/15°C night, with a 16-hour light/8-hour dark photoperiod). Leaf samples were collected at 0 (pre-inoculation), 12, 24, 48, 72, 96, and 120 hours post-inoculation (hpi), with an additional sample from PC1332 transgenics at 192 hpi. Each sample consisted of leaves from three individual plants.

Total RNA was extracted using the RNAprep Pure Plant Kit (TIANGEN, Beijing, China), and cDNA was synthesized using the MonScript™ RTIII All-in-One Mix with dsDNase (Monad, Shanghai, China). Quantitative real-time PCR (qRT-PCR) was conducted on a QuantStudio 6 Flex system (ThermoFisher, USA) using 2× SYBR Green qPCR Mix (SparkJade, Shandong, China). EF primers (Forward: 5’-TGGTGTCATCAAGCCTGGTATGGT-3’; Reverse: 5’-ACTCATGGTGCATCTCAACGGACT-3’) were used as an internal control, and LM-5 primers (Forward: 5’-GGATATCCTCATTGACCTCCGAAAATC-3’; Reverse: 5’-CCGACTTCCTAGATTGTTATTGATAGAG-3’) were used for *Yr9* amplification. Relative gene expression levels were calculated using the 2^−ΔΔCt^ method. Primer sequences for quantification are provided in the supplementary table.

### Homologs of *YrChr1B*

Transcript #1460 (*YrChr1B*) and its corresponding genomic sequence from WT LM15 have been deposited in NCBI GenBank under the accession numbers PQ412819 and PQ412818, respectively. The similarity of genes was analyzed by Blastn searches in WheatOmics (http://wheatomics.sdau.edu.cn/).

### Genetic diversity in rye and other Triticeae

Whole-genome sequencing (WGS) data for Lo7, Weining and another 13 rye inbred lines (11, 14-15) were obtained from the NCBI databases under accessions PRJNA737291, PRJEB6215, and PRJEB34439, with an average coverage of 5–6× per line. The WGS data were mapped to the Lo7 reference genome using STAR (v2.7.10a). SNP calling and filtering were carried out using GATK v4.2, with the following filtering criteria: “DP < 5 || FS > 60.0 || MQ < 40.0 || QD < 2.0”. SNPs from biallelic sites within the physical region of the *Yr9* allele, spanning from the ATG start to the TGA stop codons in the Lo7 reference genome, were extracted using Bcftools (Version 1.6) and custom Python scripts.

We further investigated the diversity of the *Yr9* gene in 19 rye germplasm lines provided by Prof. Huagang He from Jiangsu University, China. To assess the distribution of *Yr9*, two pairs of primers were employed. The first pair consisted of a forward primer (5′-GCTTGGCAACTCTGTTCTAGAG-3′) and a reverse primer (5′-CTGCATATGCCAGTAAAGGTGGT-3′). The second pair included a forward primer (5′-GTTTGTCGTTTTAAGATGCATGCCT-3′) and a reverse primer (5′-AAGGCTATTCATCTAGCTCATAATTATC-3′).

## Data Availability

All study data are included in the article and *SI Appendix*.

## Acknowledgments

We thank Prof. Huagang He for providing rye seeds. This work was financially supported by the Key Research and Development Programs of Shandong Province (2024LZGC001, 2023TZXD086).

## Author Contributions

D.F. and J.W. conceptualized the study. F.N. and J.Q. created the EMS mutant population. S.L. and J.L.L. screened *Pst-*susceptible mutants. Y.Y., S.L. and F.N. performed STAM analysis and identified the candidate gene. J.J.L., Q.C., H.S. and C.P. conducted experiments. Y.Y., J.J.L., D.F., J.W. and L.E. wrote the paper. All authors approved the final paper.

## References

1) S. Savary et al., The global burden of pathogens and pests on major food crops. Nat. Ecol. Evol. 3, 430–439 (2019).

2) S. Cheng et al., Harnessing landrace diversity empowers wheat breeding. Nature 632, 823831 (2024).

3) C. Sansaloni et al., Diversity analysis of 80,000 wheat accessions reveals consequences and opportunities of selection footprints. Nat. Commun. 11, 4572 (2020).

4) L. A. Crespo-Herrera, L. Garkava-Gustavsson, I. Ahman, A systematic review of rye (Secale cereale L.) as a source of resistance to pathogens and pests in wheat (Triticum aestivum L.). Hereditas 154, 14 (2017).

5) R. Mago et al., High-resolution mapping and mutation analysis separate the rust resistance genes Sr31, Lr26 and Yr9 on the short arm of rye chromosome 1. Theor. Appl. Genet. 112, 41–50 (2005).

6) J. Tong et al., Genome-wide atlas of rust resistance loci in wheat. Theor. Appl. Genet. 137, 179 (2024).

7) F. Wei, R. A. Wing, R. P. Wise, Genome dynamics and evolution of the Mla (powdery mildew) resistance locus in barley. Plant Cell 14, 1903–1917 (2002).

8) Z. He, S. Rajaram, Z. Xin, G. Huang. Eds. A history of wheat breeding in China. Mexico, D.F.: CIMMYT. (2001).

9) F. Ni et al., Sequencing trait-associated mutations to clone wheat rust-resistance gene YrNAM. Nat. Commun. 14, 4353 (2023).

10) X. Shi et al., Comparative genomic and transcriptomic analyses uncover the molecular basis of high nitrogen-use efficiency in the wheat cultivar Kenong 9204. Mol. Plant 15, 1440–1456 (2022).

11) M. T. Rabanus-Wallace et al., Chromosome-scale genome assembly provides insights into rye biology, evolution and agronomic potential. Nat. Genet. 53, 564–573 (2021).

12) R. Mago et al., The wheat Sr50 gene reveals rich diversity at a cereal disease resistance locus. Nat. Plants 1, 15186 (2015).

13) F. Jamal et al., Serine protease inhibitors in plants: Nature’s arsenal crafted for insect predators. Phytochem Rev. 12, 1–34 (2013).

14) E. Bauer et al., Towards a whole-genome sequence for rye (Secale cereale L.). Plant J. 89, 853–869 (2017).

15) G. Li et al., A high-quality genome assembly highlights rye genomic characteristics and agronomically important genes. Nat. Genet. 53, 574–584 (2021).

16) Y. Wang et al., An unusual tandem kinase fusion protein confers leaf rust resistance in wheat. Nat. Genetics 55, 914–920 (2023).

17) T. Jordan et al., The wheat Mla homologue TmMla1 exhibits an evolutionarily conserved function against powdery mildew in both wheat and barley. Plant J. 65, 610–621 (2011).

18) R. F. Line, A. Qayoum, Virulence, aggressiveness, evolution, and distribution of races of ‘Puccinia striiformis’ (the cause of stripe rust of wheat) in North America, 1968-87. Technical bulletin (USA) (1992), pp. 1–44.

19) F. Ni et al., Wheat Ms2 encodes for an orphan protein that confers male sterility in grass species. Nat. Commun. 8, 15121 (2017).

20) Y. Ishida, et al. Wheat (Triticum aestivum L.) transformation using immature embryos. Methods Mol. Biol. 1223, 189–198 (2015).

21) H. Liu et al., Identification of three wheat near isogenic lines originated from CB037 on tissue culture and transformation capacities. Plant Cell Tiss. Org. 152, 67–79 (2022).

